# The Regulatory Evolution of the Primate Fine-Motor System

**DOI:** 10.1101/2020.10.27.356733

**Authors:** Morgan Wirthlin, Irene M. Kaplow, Alyssa J. Lawler, Jing He, BaDoi N. Phan, Ashley R. Brown, William R. Stauffer, Andreas R. Pfenning

**Author notes:** Correspondence to (A.R.P.).

## Abstract

In mammals, fine motor control is essential for skilled behavior, and is subserved by specialized subdivisions of the primary motor cortex (M1) and other components of the brain’s motor circuitry. We profiled the epigenomic state of several components of the Rhesus macaque motor system, including subdivisions of M1 corresponding to hand and orofacial control. We compared this to open chromatin data from M1 in rat, mouse, and human. We found broad similarities as well as unique specializations in open chromatin regions (OCRs) between M1 subdivisions and other brain regions, as well as species- and lineage-specific differences reflecting their evolutionary histories. By distinguishing shared mammalian M1 OCRs from primate- and human-specific specializations, we highlight gene regulatory programs that could subserve the evolution of skilled motor behaviors such as speech and tool use.

## Main Text

Motor behavior is the primary output of the brain and a fundamental requirement for organismal survival. Fine motor control represents an elaboration upon basic movement patterns, requiring dedicated motor cortical circuitry to allow for precise, highly skilled movements (Porter and Lemon, 1993). Although the anatomical and electrophysiological mechanisms that enable motor control have become increasingly clear (Arber and Costa, 2018), the precise mechanisms linking genome sequence changes to behavioral phenotypic evolution remains a fundamental challenge in neurogenomics.

Numerous studies have explored the key contributions of individual protein-coding genes at various levels of the fine motor control circuitry. At the peripheral level, sequence and transcriptional changes in HOXC9 appear to have played a critical role in the evolution of limb- and digit-innervating spinal motor neurons in vertebrates (Jung et al., 2014). Fine motor control of limbs and digits is dependent on corticofugal neurons that project directly from motor cortex to the spinal cord (Porter and Lemon, 1993). The specific axonal projection targets of corticofugal neuron subtype are largely governed by activity of the transcription factor FEZF2 and its cofactors (Han et al., 2011; Lodato et al., 2014).

However, connecting individual genetic changes to specific fine motor phenotypes has proven challenging. Although fine motor control and tool usage in humans and chimpanzees has been shown to be highly heritable (Hopkins et al., 2015; Missitzi et al., 2013), identifying the genetic substrate for these behaviors has remained elusive. Comparative studies in humans and songbirds have identified FOXP2 as a key transcription factor for fine vocal motor control, likely through its involvement in the development and maintenance of neuroplasticity (Haesler et al., 2007; Spiteri et al., 2007). However, FOXP2 regulates the development of a wide variety of tissues beyond the brain, and knockout studies indicate that it may be associated more generally with motor skill learning (Groszer et al., 2008). Thus, there is a need for an improvement over individual gene-centric approaches that cannot account for the full complexity of the neural circuitry and electrophysiological specializations required for the evolution of fine motor behavior.

Although the majority of biological techniques and computational models for relating genetic differences to phenotypic diversity focus on genetic variation in protein-coding genes, it is widely accepted that the many of the genetic differences that influence phenotypic differences across vertebrates lie within non-coding regulatory regions, primarily enhancers (Cheng et al., 2014; King and Wilson, 1975; Pennacchio et al., 2013; Wray, 2007). Reporter assays testing both human and chimpanzee orthologs of enhancers for their ability to drive the expression of a lacZ reporter gene in mouse embryos *in vivo* have revealed that human-specific sequence changes in conserved regulatory regions can lead to tissue-specific expression in the forebrain (Kamm et al., 2013) and in the wrist and thumb (Prabhakar et al., 2008). Thus, it seems highly promising to explore how differences in gene regulatory elements could subserve the evolution of skilled motor behavior.

Recently, studies have begun to characterize the epigenomic properties of motor cortex in both rodents and primates (Adkins et al., 2020; Bakken et al., 2020; Li et al., 2020; Yao et al., 2020; Ziffra et al., 2020). However, none, to our knowledge, have attempted to explore differences within subdivisions of primary motor cortex that are associated with distinct behavioral phenotypes, and none have investigated the comparative regulatory genomic specializations of motor versus premotor cortical regions. Thus, a more refined approach is needed to identify the genomic determinants of fine motor behavior.

We set out to identify the genomic determinants of fine motor behavior, focusing on two of the most well-studied fine motor behaviors in an evolutionary neurobiological context, vocalization and manual dexterity. These behaviors are controlled by specialized brain areas and circuitry, including dedicated subdomains of the primary motor cortex (Hast et al., 1974; Rathelot and Strick, 2006, 2009; Simonyan, 2014). In order to identify the candidate gene regulatory regions uniquely active in these brain areas, we generated open chromatin data from the macaque orofacial and hand/forelimb motor cortex subdomains, additional components of the motor system in the cortex and basal ganglia, non-motor brain areas, and non-brain tissues. In order to distinguish more general regulatory genomic properties of the mammalian motor system from specializations unique to primates, we compared these macaque open chromatin regions to previously published datasets from human and mouse, and collected additional open chromatin data from rat.

We identified multiple sets of brain region-specific and species-specific open chromatin regions, including sets with conserved activity across mammals, sets with specialized activity in primates, and a set uniquely active in humans. Some of these regions are near genes that have been implicated in known aspects of motor behavior, as well as genes associated with neuromotor disabilities. These findings provide insight into the epigenomic underpinnings of fine motor control in primates, as well as providing candidate regulatory specializations that may underlie the evolution of their enhanced capacity for fine motor control.

## Results

### Generating a multi-tissue, multi-species atlas of chromatin accessibility

To identify the epigenomic specializations for fine motor behavior in the primate motor system, we isolated 11 brain areas and 2 non-brain control tissues (liver and pectoralis muscle) from two adult Rhesus macaques for ATAC-seq (Assay for Transposase-Accessible Chromatin using sequencing (Buenrostro et al., 2013) (Fig. 1, Methods). These brain areas included 2 well-characterized subdivisions of primary motor cortex (Hast et al., 1974; Rathelot and Strick, 2006, 2009; Simonyan, 2014), hand and forearm primary motor cortex (hM1), and orofacial primary motor cortex (ofM1); and 2 premotor regions, Area 6V of the ventrolateral prefrontal cortex (6V) and the supplementary motor area (SMA). As a non-frontal lobe cortical contrast, we also isolated the caudal parabelt region of the temporal lobe, corresponding to a portion of the secondary auditory cortex (2A). For non-cortical contrasts, we isolated the putamen, caudate nucleus, and nucleus accumbens (NAcc) of the striatum. From one individual, we also collected subdivisions of primary somatosensory cortex corresponding to our motor cortical subdivisions, hand (hS1) and orofacial somatosensory cortices (ofS1), as well as cerebellum.

**Figure 1.**
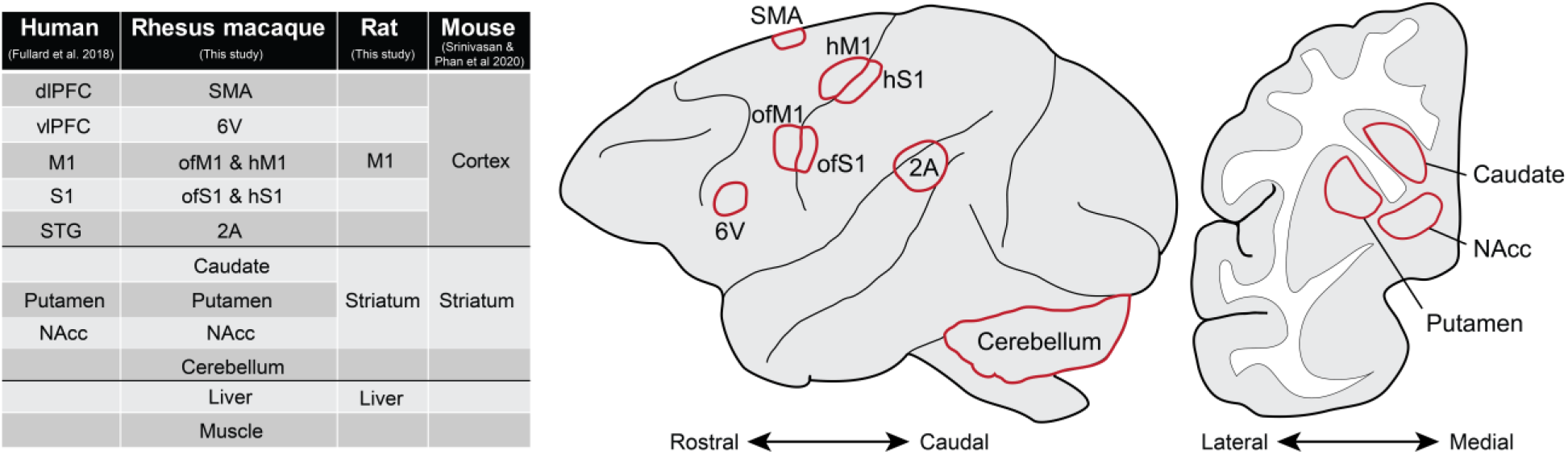
A Multi-Species Open Chromatin Atlas. Table at left presents the complete set of tissues analyzed in this study, including Rhesus macaque and rat data collected for this study as well as human (Fullard et al., 2018) and mouse (Srinivasan et al., 2020) data collected previously. Rows indicate approximate equivalence between brain areas, although we note that all macaque cortical areas are acute subregions within or proximal to the broader human regions collected. Rhesus macaque brain schematics display anatomical locations of regions processed for open chromatin data. Sagittal view (center) presents superficially visible structures while coronal view (right) presents internal structures of the basal ganglia. 2A, secondary auditory cortex; 6V, premotor area 6V; dlPFC, dorsolateral prefrontal cortex; hM1, hand and forearm M1; hS1, hand and forearm S1; M1, primary motor cortex; ofM1, orofacial M1; ofS1, orofacial S1; NAcc, nucleus accumbens; S1, primary somatosensory cortex; SMA, supplementary motor area; STG, superior temporal gyrus; vlPFC, ventrolateral prefrontal cortex.

In order to distinguish primate lineage-from species-specific specializations, we reprocessed publicly available NeuN-sorted ATAC-seq data from several human brain regions (Fullard et al., 2018) roughly comparable to some of those collected from macaque: primary motor cortex (M1), ventrolateral prefrontal cortex (vlPFC), dorsolateral prefrontal cortex (dlPFC), superior temporal gyrus (STG), NAcc, and putamen (see Fig. 1 for approximate regional equivalencies).

In order to distinguish primate-specific epigenomic specializations from more general properties of mammalian motor cortex, we also collected M1, striatum, and liver from 3 rats and processed them for ATAC-seq (Methods). In order to distinguish rodent lineage-from species-specific epigenomic features, we incorporated ATAC-seq data from cortex and striatum of 4 C57Bl/6J mice generated previously (Srinivasan et al., 2020).

For all macaque and rat samples, we prepared nuclei suspensions from fresh tissue, performed ATAC-seq as described previously (Buenrostro et al., 2013; Buenrostro et al., 2015), and sequenced the resulting libraries using Illumina NovaSeq 6000 (Methods). We used the ENCODE ATAC-seq pipeline to process all sequenced samples, integrate tissue replicates between individuals of the same species, and identify open chromatin peaks (Methods). Quality control metrics produced by this pipeline confirmed high periodicity in at least one sample per tissue from each subject, indicative of the successful preservation of the open chromatin landscape of each tissue (Fig. 2). We filtered raw open chromatin peaks to exclude coding and non-coding exonic regions as well as promoters, whose chromatin state is not primarily predictive of gene regulatory activity (Chereji et al., 2019). This filtering provided us with high-confidence sets of open chromatin region (OCR) peaks for each tissue. We applied standard motif identification and enrichment tools (McLeay and Bailey, 2010) to identify overrepresented transcription factor binding sites (TFBSs) in OCRs across tissues (Methods). The top TFBSs enriched in our brain OCR peak sets were overwhelmingly associated with known brain transcription factors—including NEUROD1, FOS::JUN, MEF2C, and TCF4—supporting our confidence that these peak sets are likely to be true gene regulatory elements in the brain regions sampled.

**Figure 2.**
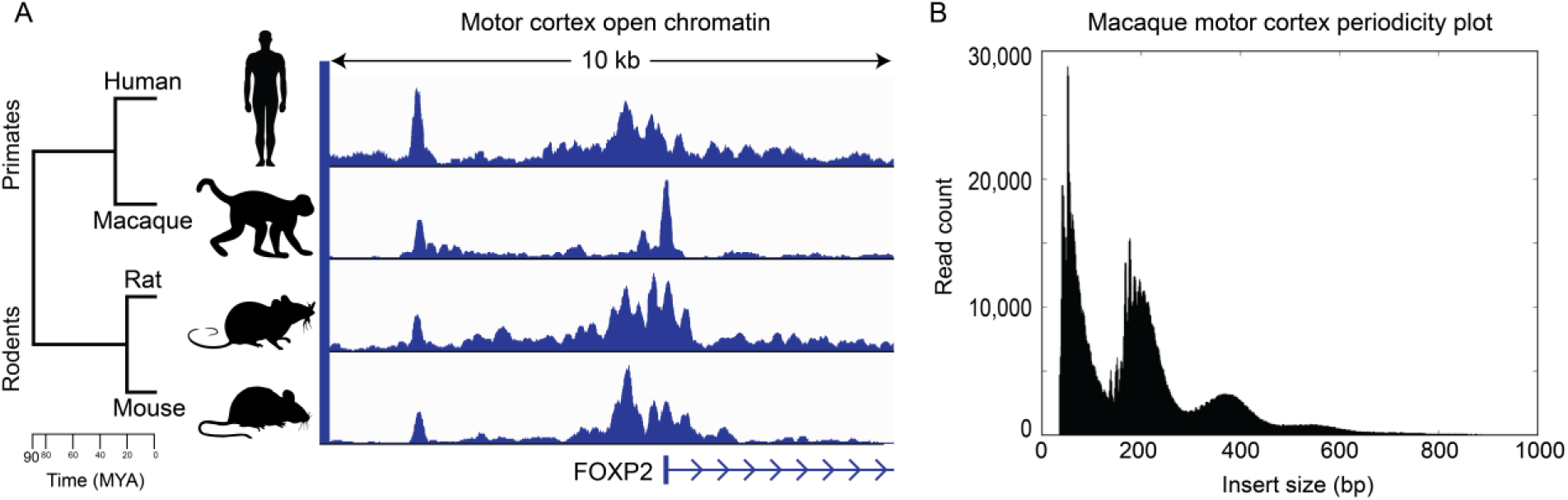
Motor Cortex Open Chromatin in Primates and Rodents. (**A**) M1 open chromatin status of human, macaque, rat, and mouse at the FOXP2 promoter. (**B**) Representative fragment length distribution of ATAC-seq libraries from macaque hM1.

### Epigenomic specializations of the non-human primate motor system

As our primary goal was to uncover potential gene regulatory specializations underlying primates’ capacity for skilled motor behavior, we sought to identify OCRs with enriched activity in specific component of the motor control system, M1 (and its subdivisions), premotor area 6V, and putamen, the primary striatal projection target of M1 (Fig. 1). Differentially active OCRs were defined as those exhibiting a log fold difference between tissue contrasts >1.5 at an adjusted p-value < 0.05 (Methods).

The number of differentially active OCRs identified between regions followed expectations given these regions’ known differences in neurobiology. Accordingly, the largest differences were observed between putamen and M1 (197,089 differentially active OCRs), reflecting the considerable functional, anatomical, and molecular differences between striatum and cortex. Considerably fewer OCRs were found to be differential between M1 and 6V (8,852 differentially active OCRs), reflecting the extensive functional similarities between these adjacent cortical regions. Between the hand and orofacial subdivisions of M1, we identified 2,225 differentially active OCRs, just 0.7% of the total number of 311,182 OCRs active in both M1 subdivision combined, highlighting the broadly conserved functional properties of M1. This suggests that the gene regulatory differences that contribute to these M1 subdivisions’ known differences in connectivity and function may be extremely subtle.

In order to interpret the biological significance of differentially active OCR sets in the primate motor system, we conducted gene ontology (GO) analysis using GREAT (McLean et al., 2010), using as a background the consensus set of reproducible OCRs across all macaque tissues (Methods).

We first sought to identify the potential biological functions of OCRs with differential activity between motor cortex subdivisions. OCRs active in orofacial M1 relative to hand M1 were associated with multiple brain-related functional terms; including myelination, neuron apoptotic processes, and forebrain neuron fate determination; as well as severe human motor disease-associated phenotypes including dysarthria, spastic paraplegia, and degeneration of the lateral corticospinal tracts (Martinez-Lage et al., 2012; Matsufuji et al., 2013; Pfeffer et al., 2015; Sacco et al., 2010; Svenstrup et al., 2011) (Table S1). Conversely, OCRs with higher activity in hand M1 were associated with functional terms related to oxidoreductase activity, smooth muscle cell regulation, and stress-activated protein kinase signaling; as well as genes associated with axon pathology and motor dysfunction (Saifetiarova and Bhat, 2019) (Table S1).

We next sought to identify epigenomic specializations that could be related to the functions of premotor and motor areas in non-human primates. OCRs active in 6V relative to whole M1 were associated with functional terms related to brain function, including interneuron migration, serotonin signaling, as well as receptor activity of peptide hormones such as somatostatin and calcitonin (Table S1). OCRs with higher activity in M1 relative to 6V were associated with functional terms related to neurotransmitter transport and secretion, synaptic vesicle processes, and voltage-gated potassium and calcium channel activity. Interestingly, M1-active OCRs were also associated with genes with known roles in dysphasia, axonal degeneration, and reduced ankle reflexes (Köhler et al., 2019) (Table S1).

Finally, we examined OCRs differentially active between M1 and putamen to identify potential functional enrichments related to these two primary motor components of the cortex and striatum, respectively. We found that OCRs selectively active in putamen were associated with expected functional processes such as dopamine receptor signaling, but also with numerous motor disease phenotypes, including ataxia, muscle weakness in upper limbs, hand tremor, and facial myokymia (involuntary twitching of the facial muscles) (Köhler et al., 2019) (Table S1). OCRs with enriched activity in M1 were associated with terms related to dendritic morphology, G-protein coupled receptor signaling, and transcriptional regulation (Table S1). Interestingly, several M1-enriched OCRs were clustered around genes associated with late-onset distal muscle weakness and progressive loss of acquired language and hand skills (Köhler et al., 2019; Vuillaume et al., 2018).

### Primate- and human-specific epigenomic specializations of the primary motor cortex

We performed a series of comparative analyses using the full set of OCRs identified in human, macaque, mouse, and rat in order to elucidate the evolution of specializations of the motor control circuit. In order to identify the set of orthologous OCRs conserved across species, we aligned open chromatin data from each tissue of each species to all other species considered in this study using a set of mammalian whole-genome sequence alignments (Armstrong et al., 2019; Hickey et al., 2013). These OCR alignments were filtered and assembled into high-confidence orthologs with a post-processing tool specifically developed for mapping regulatory elements across distantly related species (Zhang et al., 2020) (Methods). This allowed us to parsimoniously distinguish species- and lineage-specific OCRs from those that may be more generally conserved among mammals.

A majority of the 113,041 OCRs active in human M1 were found to have orthologous loci in macaque and rodents. We identified 110,908 (98.1%) orthologs of human M1 OCRs in macaque, reflecting the whole-genome sequence conservation level of 96% for this species pair (Yates et al., 2020). Likewise, in the more distantly related rat and mouse we identified orthologs for 92,031 (81.4%) and 90,935 (80.4%) of the human M1 OCR set, in line with an overall genome sequence conservation between human and both of these species of ~83% (Yates et al., 2020). However, of those OCR orthologs, far fewer showed conservation of regulatory activity. Whereas in macaque, 103,939 (93.7%) of human M1 OCR orthologs overlapped a macaque M1 OCR, in rat and mouse only 44,345 (48.8%) and 40,911 (44.5%) of aligned human M1 OCRs, respectively, overlapped a motor cortical OCR in that species. A similar pattern of high conservation of orthologous OCR loci between primate and rodents but low conservation of open chromatin state was observed in striatum, suggesting a nonlinear relationship between sequence conservation and conservation of regulatory activity.

In order to identify OCRs with primate-specific activity in M1, we identified the set of human M1 OCRs that overlapped with the previously described sets of OCRs differentially active in the macaque motor system: specifically, between subdivisions of macaque M1, those active in M1 relative to 6V, and those active in M1 relative to putamen (Table 1). We first restricted these sets to consider only those OCRs that had a clear orthologous locus in all other species (human, rat, and mouse) for which we had M1 data. We next identified the subsets of these macaque M1 OCR ortholog sets that overlapped M1 OCRs within each other species (Table 1). We found that across all contrasts, macaque M1-specialized OCRs were nearly always primate-specific, having shared activity in human M1 only (~80 - 90% of OCR orthologs, see Table 1), and very rarely displaying conserved activity in both human and either rat or mouse (~10 - 20% of OCR orthologs, see Table 1). These figures were much lower than the differences in overall conservation of M1 regulatory activity between primates (93.7%) and rodents (48.8% in rat, 44.5% in mouse). This suggests that OCRs with specialized regulatory activity in M1 relative to other regions are also less likely to be shared M1 OCRs in other species, with this likelihood of shared activity decreasing as evolutionary distance increases.

**Table 1:**
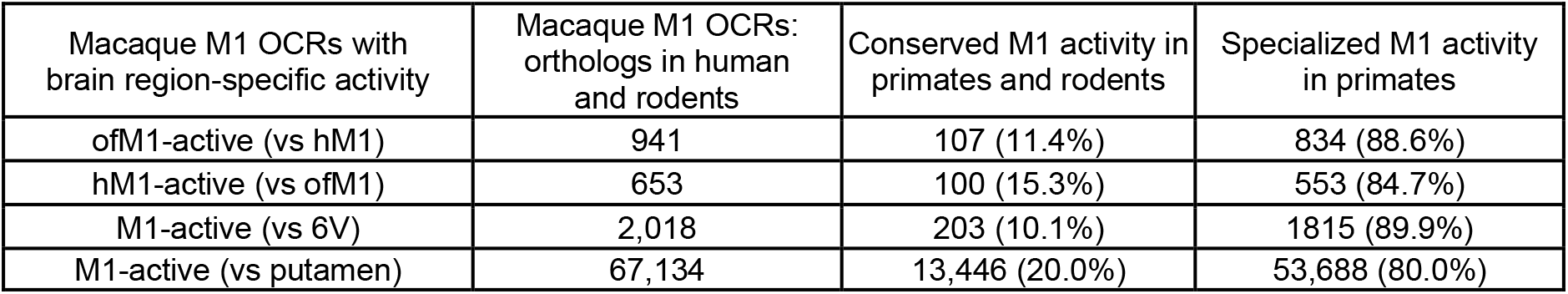
Epigenomic specializations of the primate primary motor cortex. Percentages are out of the total of macaque M1-enriched OCRs with orthologs in human and rodents. Abbreviations: OCRs: open chromatin regions; M1: primary motor cortex; ofM1: orofacial M1; hM1: hand and forelimb M1; 6V: premotor area 6V.

| Macaque M1 OCRs with brain region-specific activity | Macaque M1 OCRs: orthologs in human and rodents | Conserved M1 activity in primates and rodents | Specialized M1 activity in primates |
| --- | --- | --- | --- |
| ofM1-active (vs hM1) | 941 | 107 (11.4%) | 834 (88.6%) |
| hM1-active (vs ofM1) | 653 | 100 (15.3%) | 553 (84.7%) |
| M1-active (vs 6V) | 2,018 | 203 (10.1%) | 1815 (89.9%) |
| M1-active (vs putamen) | 67,134 | 13,446 (20.0%) | 53,688 (80.0%) |

Finally, we sought to identify the set of open chromatin regions that were uniquely specialized in human M1. We restricted our comparison to the sets of OCRs that were uniquely enriched in M1 relative to striatum (specifically putamen in human and macaque), the only brain regions for which data were available from all four species. Of the complete set of 55,732 OCRs enriched in human M1, 97% successfully mapped to macaque and ~74% mapped to rat and mouse, comparable to the overall rates of OCR alignments observed for the macaque M1 OCRs. We then removed from this set any OCRs that overlapped an OCR from any macaque, rat, or mouse brain tissue examined. This resulted in a set of 2,143 OCRs with human-specific M1 activity. Gene ontology analyses revealed this set to be associated with terms such as telomere maintenance in response to DNA damage and upregulation of histone H3K9 methylation, known to be associated with transcriptional silencing (Hyun et al., 2017) (Table S1). Human-specific M1 OCRs were also proximal to genes such as POLG and AARS2 associated with late-onset muscle weakness, motor neuropathy, and dysarthria (Köhler et al., 2019; Lynch et al., 2016; Van Goethem et al., 2003) (Table S1).

Within the set of human-specific M1 OCRs, we identified a region downstream of FOXP2, a gene well characterized for its role in speech disability (Lai et al., 2001; White et al., 2006). To our knowledge, this particular element (chr7:114,819,495 - 114,820,276, hg38) has never been reported in the experimental literature on FOXP2 regulatory genomics (Atkinson et al., 2018; Becker et al., 2015; Caporale et al., 2019; Maricic et al., 2013; Moralli et al., 2015; Turner et al., 2013). This locus is, however, proximal to a noncoding region that has previously been associated with childhood apraxia of speech when interrupted through a natural chromosomal inversion event (Moralli et al., 2015). Although this OCR was detected on the basis of being differentially active in M1 relative to putamen, we note that within the additional human brain open chromatin datasets examined in this study, appreciable levels of activity in vlPFC, dlPFC, and STG were also detected; suggesting that its functions may extend into other cortical domains beyond M1 as well.

## Discussion

With the goal of identifying the gene regulatory properties of the primate fine motor system, we profiled the genome-wide open chromatin state of 13 macaque and 3 rat tissues, including behaviorally relevant subdivisions of macaque M1 whose open chromatin state had not previously been assessed. In so doing, we have generated a high-quality epigenomic resource to facilitate further discoveries on the regulatory biology of the brain in these important model systems. The open chromatin regions (OCRs) identified from these experiments are overwhelmingly associated with genes and enriched for TFBSs with known roles in the brain, suggesting that they are in fact likely to represent gene regulatory enhancers. This possibility could be further explored through the integration of transcriptome data from comparable subdivisions of the motor system, which would facilitate associating regionally active OCRs with the differentially expressed genes they may potentially regulate.

We identified open chromatin specializations unique to specific components of the non-human primate motor system. We found motor system-enriched OCRs to be clustered around genes that are involved in functional processes relevant to brain function, including neurotransmitter transport, synaptic vesicle processes, and voltage-gated ion channel activity. Between the orofacial and hand M1 subdivisions, several such genes were related to oligodendrocyte-mediated myelination processes, which are known to be specifically relevant to motor learning in M1 (Scala et al., 2020). Gene regulatory elements under ongoing evolutionary selection in the hominin lineage have recently been demonstrated to be primarily associated with oligodendrocyte function (Castelijns et al., 2020), suggesting that these processes may be a critical component in the evolution of the primate fine motor system.

Many of these genes are also known to be associated with severe motor disability. Among the top genes associated with OCRs differentially regulated between orofacial and hand M1 were SPG7, NIPA1, and PLP1. In humans, mutations in SPG7 are a common cause of hereditary spastic paraplegia, dysarthria, and other forms of ataxia in humans (Pfeffer et al., 2015), as well as in KO mice, where the gene has been confirmed to be expressed in neocortical pyramidal cells (Sacco et al., 2010). NIPA1 is similarly associated with spastic paraplegia, dysarthria, and atrophy of the small hand muscles in humans (Svenstrup et al., 2011), as well as widespread pyramidal motor neuron loss in the motor cortex and other areas (Martinez-Lage et al., 2012). Functional loss of PLP1 is associated with a severe form of hereditary spastic paraplegia known as Pelizaeus–Merzbacher disease, which is characterized by significant dysarthria, ataxia, and other motor pathologies (Matsufuji et al., 2013). These links to known functions and disorders bolster confidence in the biological validity of our OCR sets, while also providing possible genomic mechanisms behind the neurobiological bases for both normal motor cortical function as well as motor disability.

We identified a number of OCRs with enriched activity in the motor cortex of human and macaque relative to rat and mouse, which could reflect gene regulatory specializations supporting the evolution of the enhanced fine motor control capabilities of primates relative to rodents. We observed that the proportion of orthologous OCRs shared between primates and rodents strongly matched their overall rates of genomic sequence conservation. However, the conservation of tissue-specific OCR activity was much lower. The percentage of OCRs with shared regulatory activity between primates and rodents was especially low in the case of OCRs that were differentially active between distinct components of the motor system. These findings are reflective of the stark disconnect between conservation of a regulatory element’s orthologous locus and conservation of regulatory activity on evolutionary timescales. The fact that the conservation OCR’s orthologous locus is not directly predictive of its tissue-specific activity suggests that there may be particular features within these sequences that are critical for orchestrating tissue-specific regulatory activity. This growing consensus is motivating a diversity of attempts to incorporate evolutionary information into machine learning models to predict these higher-order sequence features in order to elucidate the basic grammar of transcriptional regulation (Chen et al., 2018; Kelley, 2020; Minnoye et al., 2020).

We also identified a set that of OCRs with enriched activity in M1 of humans but no activity in any of the other macaque, rat, or mouse brain tissue examined in this study. As observed with OCRs with differential activity between motor cortical subdivisions, a number of these human-specific, M1-enriched OCRs were associated with genes involved in various disabilities relating to motor function. These include POLG, which has been connected to a range of ataxic neuropathies frequently characterized by dysarthria (Van Goethem et al., 2003), as well as AARS2, mutations of which are associated with adult-onset leukodystrophy characterized by motor polyneuropathy including dysarthria (Lynch et al., 2016). We also identified an OCR associated with known speech disorder gene FOXP2, which is proximal to a region where chromosomal rearrangement has been shown to result in severe childhood apraxia of speech (Moralli et al., 2015). This finding highlights one way in which adding evolutionary context can reveal insights hidden within existing human data.

We note that our candidate human-unique, M1-enriched OCRs represent specializations of M1 relative to striatum, and of human relative to macaque. Although this reveals one aspect of how the human motor system has specialized in comparison to the species and brain regions available to us in this study, it is possible that some of these specializations may reflect more general hominid specializations of the cortex or frontal lobe. Distinguishing between these possibilities will be facilitated by the availability of open chromatin datasets from a broader range of comparable brain areas from other species, as well as improved machine learning models that can predict regulatory activity from genomic sequence alone. We anticipate that cross-species, multi-tissue epigenomic data resources like those generated in the present study will facilitate the training and improvement of such models.

## Methods

### Animals and sample collection

All animal procedures were in accordance with the National Institutes of Health Guide for the Care and Use of Laboratory Animals and approved by the Institutional Animal Care and Use Committees (IACUC) of Carnegie Mellon University (Protocol ID 201600003) and the University of Pittsburgh (Protocol ID 19024431). Rhesus macaques were single- or pair-housed at the University of Pittsburgh with a 12h-12h light-dark cycle. Macaques sampled in this study were a 12-year-old female (8.1 kg) and a 4-year-old male (6.0 kg). Before surgery, macaques were initially sedated with ketamine (15 mg/kg IM), and then ventilated and further anesthetized with isoflurane. The animals were transported to a surgery suite and placed in a stereotaxic frame (Kopf Instruments). We removed the calvarium and then perfused the circulatory systems with 3-4 liters of ice cold, oxygenated macaque artificial cerebrospinal fluid (124 mM NaCl, 5 mM KCl, 2 mM MgSO_4_, 2 mM CaCl_2_, 3 mM NaH_2_PO_4_, 23 mM NaHCO_3_, 10 mM glucose). We then opened the dura and removed the brain. All brain regions were excised under a dissection microscope. To supplement adult mouse brain data collected previously (Srinivasan et al., 2020), we also collected M1, striatum, and liver tissues from three rats (1 male Sprague-Dawley, housed in the University of Pittsburgh; 2 Brown Norway, 1 male and 1 female, housed at Carnegie Mellon University). Rats were euthanized by isoflurane overdose followed by decapitation. Liver was collected immediately. Brains were sliced into 300 μm sections in a vibrating microtome (Leica VT 1200) in ice-cold, oxygenated rodent artificial cerebrospinal fluid (119 mM NaCl, 2.5 mM KCl, 1 mM NaH_2_PO_4_ (monobasic), 26.2 mM NaHCO_3_, 11 mM glucose). Brain regions of interest were sampled from these coronal sections under a dissection microscope and transferred to chilled lysis buffer (Buenrostro et al., 2015).

### ATAC-seq

Tissue samples were processed as described previously (Buenrostro et al., 2013; Buenrostro et al., 2015), with the following minor differences in procedure and reagents. Nuclei were isolated from dissected tissues using 30 strokes of homogenization with the loose pestle (0.005 in. clearance) in 5mL of cold lysis buffer placed in a 15 mL glass Dounce homogenizer (Pyrex #7722-15). The nuclei suspensions were filtered through a 70 μm cell strainer, pelleted by centrifugation at 2,000 x *g* for 10 minutes, resuspended in water, and filtered a final time through a 40 μm cell strainer. Sample aliquots were stained with DAPI (Invitrogen #D1206), and nuclei concentrations were quantified using a manual hemocytometer under a fluorescent microscope. Approximately 50,000 nuclei were input into a 50 μL ATAC-seq tagmentation reaction as described previously (Buenrostro et al., 2013; Buenrostro et al., 2015). The resulting libraries were amplified to 1/3 qPCR saturation, and fragment length distributions estimated by the Agilent TapeStation System showed high quality ATAC-seq periodicity. We shallowly sequenced barcoded ATAC-seq libraries at 1-5 million reads per sample on an Illumina MiSeq and processed individual samples through the ENCODE pipeline (Landt et al., 2012) for initial quality control. We used the QC measures from the pipeline (clear periodicity, library complexity, and minimal bottlenecking) to filter out low-quality samples and re-pooled a balanced library for paired-end deep sequencing on an Illumina NovaSeq 6000 System through Novogene services to target >30 million uniquely mapped fragments per sample after mitochondrial DNA and PCR duplicate removal.

### Data Analysis

We processed raw FASTQ files of ATAC-seq experiments with the ENCODE ATAC-seq pipeline (Landt et al., 2012) accessed at https://github.com/ENCODE-DCC/atac-seq-pipeline. To supplement our macaque and rat data, we obtained mouse brain ATAC-seq data for cortex and striatum from (Srinivasan et al., 2020). We also processed publicly available NeuN-sorted ATAC-seq data from human postmortem brain (Fullard et al., 2018) from regions roughly corresponding to those collected from macaque, namely: primary motor cortex (PMC), ventrolateral prefrontal cortex (VLPFC), dorsolateral prefrontal cortex (dlPFC), superior temporal gyrus (STC), nucleus accumbens (NAc), and putamen (PUT). We downloaded these data from the Sequence Read Archive (SRA) through Gene Expression Omnibus (GEO) accession ID GSE96949.

We ran the ENCODE pipeline using the rheMac8 assembly for macaque, the hg38 assembly for human, the rn6 assembly for rat, and the mm10 assembly for mouse. We ran the pipeline with the default parameters except for “atac.multimapping” : 0, “atac.cap_num_peak”: 300,000, “atac.smooth_win”: 150, “atac.enable_idr”: true, and “atac.idr_thresh”: 0.1. We combined technical replicates when processing data. We generated filtered bam files, peak files, and signal tracks for each replicate and the pool of replicates for each tissue, per species. We removed samples that had low periodicity indicated by ENCODE quality control metrics and reprocessed the remaining replicates. Since our replicates often differed substantially in sequencing depth, we defined reproducible peaks to be peaks with an irreproducible discovery rate (IDR, (Li et al., 2011)) < 0.1 across pooled pseudo-replicates, and used these peaks for all downstream analyses. In the case of tissues for which there was only one high-quality biological replicate, we used peaks that were reproducible according to IDR < 0.1 across self-pseudo-replicates.

In addition to identifying peak sets for individual tissues, for each species, we identified one set of peaks to serve as a genome-wide background set representing the union of the reproducible open chromatin peaks identified from all processed tissues. This background set was obtained using bedtools (Quinlan and Hall, 2010) intersect with the -wa and -u options to combine all reproducible peak sets per species. A number of steps were taken to prepare OCR peak sets for downstream analyses. Peaks within 50 bp of one another were combined using bedtools merge. We used bedtools subtract with option -A to remove those peaks which were within 2 kb from any annotated coding or noncoding exons, enabling us to exclude promoters, coding sequences, and noncoding RNAs from our background set. Peaks aligned to chromosome Y were removed to control for sex-biased effects. In order to identify the complete set of exonic exclusion regions for macaque, we used the complete set of rheMac8 RefSeq annotations (O’Leary et al., 2016) supplemented with UCSC’s ‘xenoRefSeq’ annotations obtained from their track browser, which represent RefSeq annotations from dozens of other species aligned to macaque using liftOver (Karolchik et al., 2004). For human, rat, and mouse; we used the hg38, rn6, and mm10 RefSeq annotation sets, respectively (O’Leary et al., 2016).

To identify OCR peaks differentially active between tissue contrasts, we first quantified the number of reads from each tissue aligning to that species’ consensus peakset using featureCounts (Liao et al., 2014). We then contrasted readcounts at each peak between tissues using the negative binomial model in the DESeq2 R package (Love et al., 2014). We considered peaks differential that exhibited a log fold difference between tissue contrasts >1.5 with an adjusted p-value < 0.05.

In order to identify orthologous OCRs across species, we aligned open chromatin data from each tissue of each species to all other species considered in this study. OCRs were mapped between species using halLiftover (Hickey et al., 2013) with default parameters, using the Zoonomia Cactus multiple whole-genome sequence alignment for graph-based genome coordinate mapping (Armstrong et al., 2019). The raw outputs of halLiftover were then filtered and assembled into contiguous OCRs using the halLiftover Post-processing for the Evolution of Regulatory Elements (HALPER) tool (Zhang et al., 2020), with parameters -max_frac 1.2, - min_len 50, and -protect_dist 5.

Gene ontology analyses were performed using the Genomic Regions Enrichment of Annotations Tool (GREAT) version 4.0.4 (McLean et al., 2010). Genomic coordinates of differential OCR peak sets were used as foreground regions. For the background regions, we used the union of all reproducible open chromatin peaks identified from all processed tissues per species. Significantly overrepresented ontology categories were ranked by the hypergeometric false discovery rate q-value and only GO terms made up of at least 5 genes were considered.

To identify transcription factor binding motifs enriched in differential OCR peak sets of interest relative to shuffled sequences, we used AME in MEME suite (Bailey et al., 2009; McLeay and Bailey, 2010), performing a Fisher’s exact test on the total odds score (the sum of the position weight matrix (PWM) motif scores of the sequence) with all other parameters set to default. For our PWM set, we used the JASPAR2018 CORE set of non-redundant vertebrate motifs (Khan et al., 2018).

## Author contributions

M.E.W., A.R.P., and W.R.S. designed the study. M.E.W. collected the ATAC-seq data with assistance from A.J.L., J.H., and A.R.B. M.E.W. analyzed the data with assistance from I.M.K., A.R.P., and B.N.P. M.E.W. wrote the manuscript, with feedback from all authors.

## Acknowledgements

We would like to thank Christina M. Cerkevich for her guidance in identifying macaque neuroanatomical subdivisions prior to dissection. This work used the Extreme Science and Engineering Discovery Environment (XSEDE), through the Pittsburgh Supercomputing Center Bridges Compute Cluster, which is supported by National Science Foundation grant number TG-MCB190067.

